# CVnCoV protects human ACE2 transgenic mice from ancestral B BavPat1 and emerging B.1.351 SARS-CoV-2

**DOI:** 10.1101/2021.03.22.435960

**Authors:** Donata Hoffmann, Björn Corleis, Susanne Rauch, Nicole Roth, Janine Mühe, Nico Joel Halwe, Lorenz Ulrich, Charlie Fricke, Jacob Schön, Anna Kraft, Angele Breithaupt, Kerstin Wernike, Anna Michelitsch, Franziska Sick, Claudia Wylezich, Stefan O. Müller, Thomas C. Mettenleiter, Benjamin Petsch, Anca Dorhoi, Martin Beer

**Affiliations:** Institute of Diagnostic Virology, Friedrich-Loeffler-Institut, Greifswald-Insel Riems, Germany; Institute of Immunology, Friedrich-Loeffler-Institut, Greifswald-Insel Riems, Germany; CureVac AG, Tübingen, Germany; Department of Experimental Animal Facilities and Biorisk Management, Friedrich-Loeffler-Institut, Greifswald-Insel Riems, Germany; Friedrich-Loeffler-Institut, Federal Research Institute for Animal Health, Greifswald-Insel Riems, Germany

**Author notes:** These authors contributed equally to this work (Donata Hoffmann, Björn Corleis). Corresponding authors. These authors contributed equally to this work.

## Abstract

The ongoing severe acute respiratory syndrome coronavirus-2 (SARS-CoV-2) pandemic necessitates the fast development of vaccines as the primary control option. Recently, viral mutants termed “variants of concern” (VOC) have emerged with the potential to escape host immunity. VOC B.1.351 was first discovered in South Africa in late 2020, and causes global concern due to poor neutralization with propensity to evade preexisting immunity from ancestral strains. We tested the efficacy of a spike encoding mRNA vaccine (CVnCoV) against the ancestral strain BavPat1 and the novel VOC B.1.351 in a K18-hACE2 transgenic mouse model. Naive mice and mice immunized with formalin-inactivated SARS-CoV-2 preparation were used as controls. mRNA-immunized mice developed elevated SARS-CoV-2 RBD-specific antibody as well as neutralization titers against the ancestral strain BavPat1. Neutralization titers against VOC B.1.351 were readily detectable but significantly reduced compared to BavPat1. VOC B.1.351-infected control animals experienced a delayed course of disease, yet nearly all SARS-CoV-2 challenged naïve mice succumbed with virus dissemination and high viral loads. CVnCoV vaccine completely protected the animals from disease and mortality caused by either viral strain. Moreover, SARS-CoV-2 was not detected in oral swabs, lung, or brain in these groups. Only partial protection was observed in mice receiving the formalin-inactivated virus preparation. Despite lower neutralizing antibody titers compared to the ancestral strain BavPat1, CVnCoV shows complete disease protection against the novel VOC B.1.351 in our studies.

## Introduction

Coronavirus disease 2019 (COVID-19) severely affects human health and societies worldwide. It has accounted for more than 116 million morbidities and 2.5 million fatalities by early March 2021 (WHO, https://covid19.who.int). The responsible pathogen, severe acute respiratory syndrome coronavirus type 2 (SARS-CoV-2), has rapidly spread globally despite stringent intervention strategies (*1*). To control pandemic spread and disease, vaccination is considered the most important and effective control measure (*2*). Several vaccines based on mRNA technology or viral vectors are now authorized for emergency use and further products are in final licensing phases (*3*). SARS-CoV-2 underwent adaptive mutations early during the pandemic, with the D614G variant becoming globally dominant at the beginning of 2020 (*4–6*). Viral evolution is a highly dynamic process that results in emergence of multiple, geographically distinct new variants, first identified in the UK (B1.1.7), South Africa (B.1.351) and Brazil (B.1.1.28; P1) (https://www.ecdc.europa.eu/en/publications-data/covid-19-risk-assessment-variants-vaccine-fourteenth-update-february-2021). These “variants of concern (VOC)” acquired numerous mutations, particularly in the spike protein encoding gene (S), most frequently within the S1 and the receptor binding domain (RBD) (*7–9*). These mutations confer higher binding affinities and allow some VOC to evade pre-existing immunity (*10*), resulting in increased transmissibility, including epidemiologic scenarios where “herd immunity” was expected (*11*). Whereas variant B.1.1.7 might still be efficiently neutralized by vaccination-elicited antibodies despite the RBD mutations (*12–14*), variant B.1.351 showed a remarkable resistance to sera from vaccinated, as well as from convalescent individuals (*15–18*). VOCs that evade from efficient cross-neutralization may evolve into dominant strains and necessitate vaccine efficacy re-assessment. While indications for cross neutralization exist, e.g. by sera from individuals vaccinated with SARS-CoV-2 mRNA vaccines (*13, 19*), *in vivo* data from experimental immunization/challenge studies in standardized animal models are pending. In a parallel approach, we investigated the efficacy of mRNA vaccine CVnCoV against SARS-CoV-2 using an early B lineage 614G strain and the novel VOC B1.351 in a human ACE2 transgenic mouse model of severe COVID-19 (*20*).

## Results

### Strong antibody responses in mRNA-vaccinated mice

We used the K18-hACE2 transgenic mouse model (*21*) to determine the protective efficacy of a spike-protein encoding mRNA vaccine (CVnCoV) against the ancestral SARS-CoV-2 B-lineage strain BavPat1 that closely matches the mRNA encoded S protein, and the heterologous VOC B.1.351 NW-RKI-I-0028. For immunization, either 8μg of the CVnCoV vaccine (*22*) or 20μL of a formalin-inactivated and adjuvanted SARS-CoV-2-preparation (FI-Virus) was administered on days 0 and 28. Mice were challenged 4 weeks after boost vaccination with more than 10^5^ TCID50 of SARS-CoV-2 BavPat1 or B.1.351 NW-RKI-I-0028. A sham (NaCl) group served as non-vaccinated control (Fig. S1). Sera from CVnCoV-vaccinated mice collected on days 28 and 55 showed a strong induction of anti-RBD total immunoglobulin (Ig) with a significant increase on day 55 (4 weeks after boost) compared to day 28 (4 weeks after prime). CVnCoV-induced anti-RBD total Ig levels were significantly higher than levels induced by the FI-Virus preparation. There was no significant boost effect with the FI-Virus preparation when comparing anti-RBD levels at day 55 versus day 28 (Fig. 1A). The strong induction of anti-RBD antibodies in the CVnCoV group was reflected by high virus neutralization titers (VNT). Sera from day 55 after immunization with CVnCoV showed a significantly higher neutralizing capacity compared to sera from animals that had received the FI-Virus preparation. Importantly, neutralization of VOC B.1.351 (mean = 525) was less effective compared to BavPat1 (mean = 10,151) for the CVnCoV group, but far exceeded the values recorded for the FI-Virus group (mean ≤ 32) (Fig. 1B). Overall, CVnCoV induced robust antibody responses in a prime-boost regime, capable of efficiently neutralizing both BavPat1 and VOC B.1.351 NW-RKI-I-0028 *in vitro.*

**Fig. 1.**
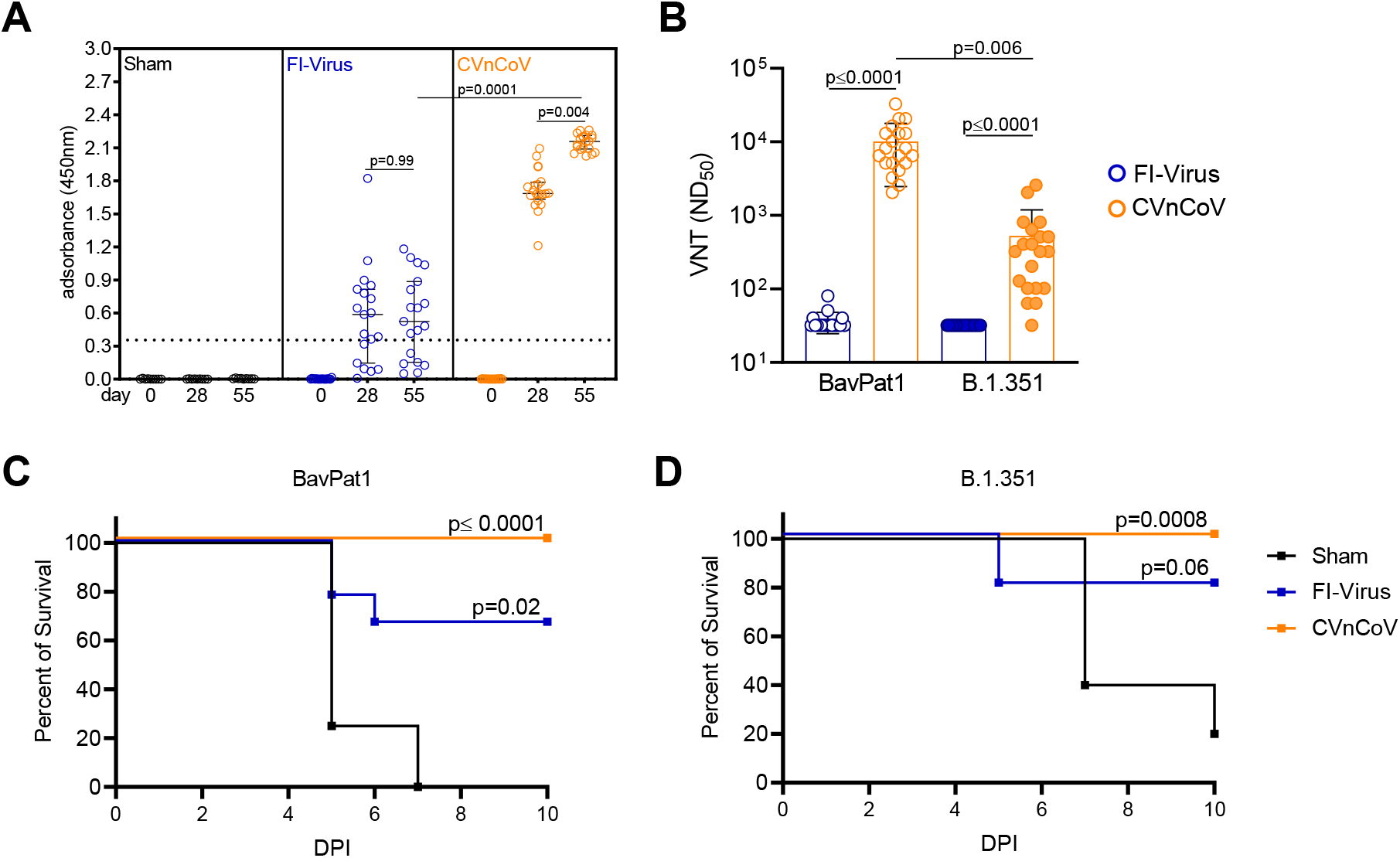
CVnCoV protects K18-hACE2 mice against SARS-CoV-2 variants BavPat1 and B1.351. K18-hACE2 mice were vaccinated with 8μg CVnCoV, received 10^6^ FI-Virus or NaCl (SHAM) on day 0 and day 28 followed by i.n. challenge with 10^5,875^ TCID50 of SARS-CoV-2 variants BavPat1 or 10^5,5^ TCID50 B1.351. (**A**) RBD-Elisa with sera from K18-hACE2 mice on day 0, 28 and 55 of respective groups: median and interquartile range are presented. Dashed line indicates threshold for positive anti-RBD antibody level. (**B**) Virus neutralization assay using day 55 sera from all three groups. Bars indicate mean with SD. (**C** and **D**) Survival curves (Kaplan-Meyer) for K18-hACE2 mice from all three groups challenged either with BavPat1 (**C**) or B.1.351 (**D**) and followed up for 10 days post infection (DPI). P-values were determined by nonparametric one-way ANOVA and Dunn’s multiple comparisons test (**A** and **B**) or log-rank (Mantel-Cox) test (**C** and **D**).

### Complete protection of mRNA-immunized mice from disease and mortality

Subsequently, the potential of CVnCoV to protect from SARS-CoV-2 challenge infection was analyzed. Stocks of both challenge viruses were characterized by deep-sequencing demonstrating the characteristic mutations of VOC B.1.351, but no other relevant alterations (Fig. S2, table S2). Immunized K18-hACE2 mice were studied using a high dose-challenge model which induces severe clinical disease resembling COVID-19 in humans (*23*). In addition, mice develop severe encephalitis specific to this animal model (*20*). On day 4, animals in the sham group started succumbing to the BavPat1 infection (Fig. 1C). B.1.351 infection led to a delayed onset of severe disease compared to BavPat1, with 20% survival on day 10 after inoculation (Fig. 1D). Thus, K18-hACE2 mice were highly susceptible to both SARS-CoV-2 variants. Importantly, vaccination with CVnCoV resulted in complete protection (100% survival) against BavPat1 and B.1.351 with no significant weight loss or disease symptoms throughout the course of the challenge infection (Fig. 1C-D and Fig. S3). In contrast, prior administration of the FI-Virus preparation provided sub-optimal protection against either BavPat1 or B.1.351, resulting in weight loss and signs of distress (Fig.1 and Fig. S3). Some of the FI-virus-immunized animals experienced very early weight loss and disease signs after VOC B.1.351 challenge infection, earlier than sham-groups. In conclusion, survival rates, body weight changes and disease scores revealed complete protection by the CVnCoV vaccine in K18-hACE2 mice against lethal SARS-CoV-2 challenge, including against VOC B.1.351.

### mRNA-immunization significantly decreased viral RNA loads in selected tissues

To investigate whether CVnCoV vaccination prevented productive infection or dissemination of replicating SARS-CoV-2, we took oral swabs on 4 dpi to monitor viral RNA load in saliva. In the sham group, 4/4 and 4/5 samples were positive for viral genome after infection with BavPat1 or VOC B.1.351, respectively (Fig. 2A). FI-Virus administration prior to challenge did not significantly reduce viral genome load in saliva with 40 to 60% of animals showing positive RT-qPCR results on 4 dpi (Fig. 2A). In contrast, after CVnCoV vaccination, no viral genomes were detected in oral swabs of either challenge group. To further explore the prevention of viral replication following challenge, we determined viral load in the upper respiratory tract (URT) (conchae) and the lower respiratory tract (LRT) (trachea, caudal lung and cranial lung), as well as in the central nervous system (brain, cerebellum/cerebrum) (Fig. 2B-F) in animals reaching the humane endpoint or at the day of termination (10dpi). Similar to the quantitative RNA load results obtained from the oral swabs, the URT provided a niche for replication in both sham and FI-Virus groups (Fig. 2B). In the CVnCoV-vaccinated group challenged with BavPat1, only 3/10 animals showed low genome copy numbers in the conchae. No animal was positive in the LRT or the brain, indicating complete protection from infection by BavPat1 (Fig. 2C-F). For VOC B.1.351, 6/10 CVnCoV-vaccinated animals exhibited residual viral replication in the conchae, but viral levels were reduced without reaching statistical significance (Fig. 2B). In contrast, CVnCoV prevented any detectable replication of this VOC in the LRT and the brain, with low viral copy numbers close to the limit of detection in the lung of only 2/10 animals, and in the cerebrum for only 1/10 animals (Fig. 2C-F). FI-Virus administration provided partial protection in the LRT in animals challenged with BavPat1, but not with VOC B.1.351, and did not significantly protect against viral replication in the cerebellum or cerebrum regardless of SARS-CoV-2 variant (Fig. 2). Of note, some of the animals receiving the FI-Virus preparation showed viral loads at the level of the sham group in the LRT (Fig. 2).

**Fig. 2.**
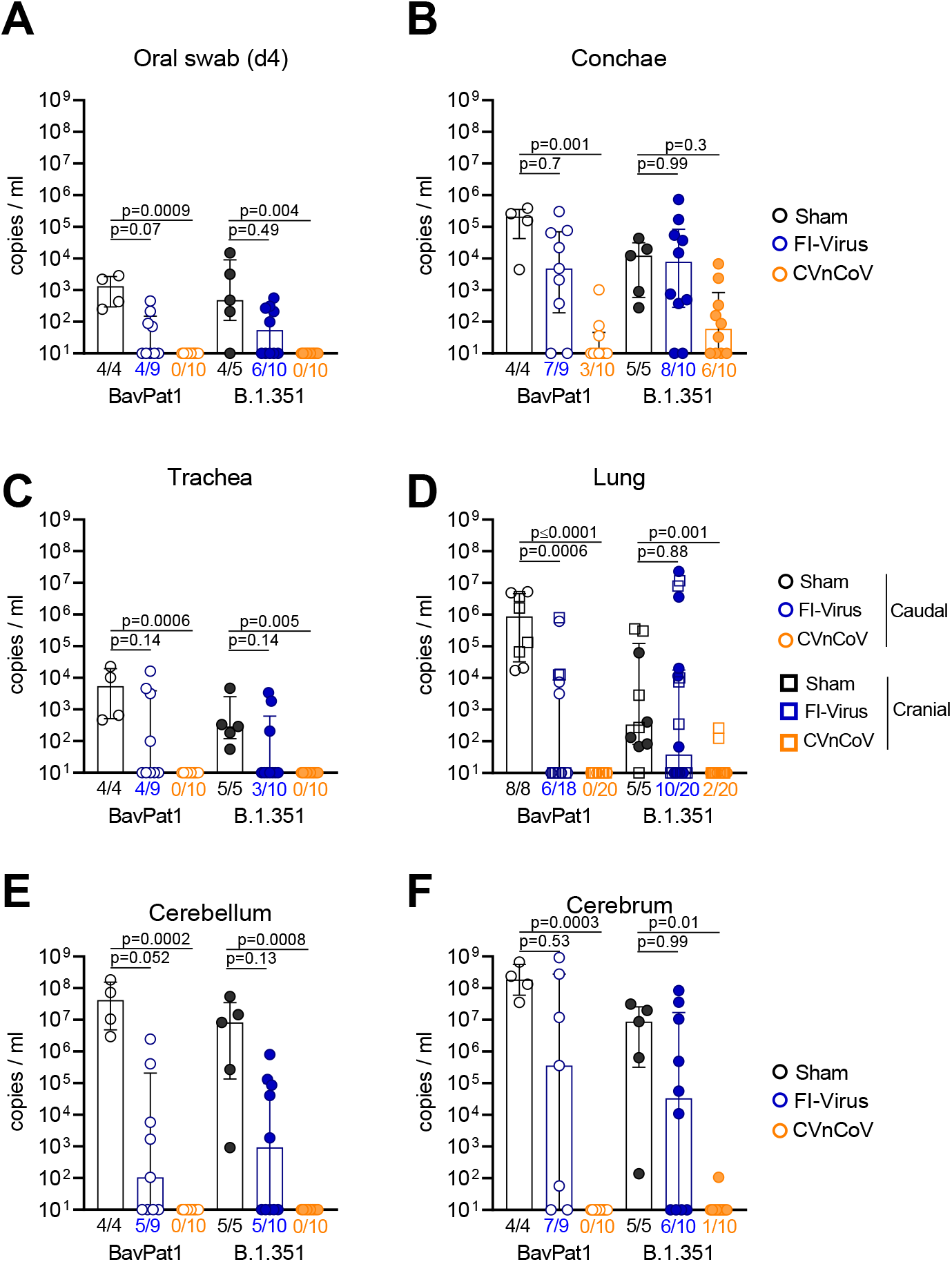
CVnCoV prevents replication of SARS-CoV-2 variants BavPat1 and B.1.351 in K18-hACE2 mice. RT-qPCR for genomic RNA of SARS-CoV-2 was performed with either (**A**) oral swab samples at day 4 or (**B**) from organ samples of the upper respiratory tract, (**C** - **E**) the lower respiratory tract, and (**F** and **G**) the brain at day 10 or at the humane endpoint. P-values were determined by nonparametric one-way ANOVA and Dunn’s multiple comparisons test. Scatter plots are labeled with median (height of the bar) and interquartile range.

In summary, vaccination with CVnCoV conferred complete protection against lethal challenge with SARS-CoV-2 lineage B BavPat1 and VOC B.1.351 strain NW-RKI-I-0028 in the transgenic K18-hACE2 mouse model. CVnCoV induced robust anti-RBD antibody responses with high neutralizing capacity against both BavPat1 and VOC B.1.351. Furthermore, CVnCoV prevented dissemination of SARS-CoV-2 from the inoculation site into other organs and provided a solid protection against an ancestral SARS-CoV-2 and a VOC B.1.351 strain.

## Discussion

The emergence of new strains with immune escape potential, such as the VOC of the B.1.351 lineage that appeared first in South Africa, are of great concern, since all available COVID-19 vaccines are based on the ancestral SARS-CoV-2 strains. The B.1.351 variant is of particular interest due to the observed immune-escape features with a reduced neutralization efficacy (*10*) and reduced protective efficacy reported for a licensed vaccine (*24*). We therefore tested an mRNA vaccine (CVnCoV) against a standard ancestral SARS-CoV-2 B lineage strain (BavPat1) in comparison to a VOC B.1.351 isolate in a transgenic mouse model.

Our data demonstrate that CVnCoV fully protects mice against disease caused by two different SARS-CoV-2 variants. CVnCoV vaccination, but not immunization with FI-Virus, rescued all transgenic mice from lethal infection caused by BavPat1 and VOC B.1.351 isolate NW-RKI-I-0028. The sub-optimal FI-Virus preparation reduced viral replication in the LRT solely after challenge with BavPat1, but showed no significant effect on viral dissemination as well as the viral genome loads in the URT. In contrast, CVnCoV immunization resulted in abundant RBD-specific and neutralizing antibodies, and conferred a complete and robust protection, including from viral replication in the lung and the brain. Only very limited viral replication was observed in the URT of mRNA-vaccinated animals challenged with VOC B.1.351. The relevance to disease transmission of this minimal viral replication in the conchae remains to be established.

The reduced neutralizing capacity of sera from CVnCoV-vaccinated transgenic mice against VOC B.1.351, and the insufficient prevention of replication in the conchae, might reflect the currently detected transmission rates of this VOC in human populations previously exposed to the ancestral strain. Nevertheless, our study provides the first evidence for the efficacy of a vaccine to prevent disease and viral dissemination from the site of infection against an emerging SARS-CoV-2 variant in a sensitive, well-established and accepted *in vivo* model. The very high neutralizing titers against BavPat1 elicited by the mRNA immunization as well as the fold reduction recorded for VOC B.1.351 may be unique to the mouse model employed in this study and require validation in other experimental models.

The pathophysiology of SARS-CoV-2 VOC infections remains largely unknown and detailed animal model data are missing. In our study, we observed a delayed course of disease in K18-hACE2 mice infected with a VOC B.1.351 strain, and hypothesize that mutation accumulation might result in a changed *in vivo* phenotype. Short-term infections performed in hamsters have failed to detect diverging phenotypes in ancestral versus VOC lineages (*25*), but comparable data sets about the complete course and replication in the upper respiratory tract are still pending for all VOCs in other animal models. These apparently discordant findings call for further pathological and immunological assessments of pathogenicity of emerging lineages.

Here we report full protection against a VOC by CVnCoV immunization, associated with high anti-RBD and neutralizing antibodies. Whether antibodies alone were sufficient for the beneficial outcome remains to be further validated. Broad immune responses elicited by vaccines, including cellular responses in addition to neutralizing antibodies, antibody dependent cytotoxicity (ADCC), or antibody-mediated innate immune effector functions, could help explain the protection. Potent T cell immunity could ensure the success of an immunization when antibodies decline. CVnCoV vaccination was previously shown to induce Th1 immunity and trigger S-specific CD8^+^ T cell responses in mice (*22*). Re-challenge studies in non-human primates confirmed a role for CD8^+^ T cells in protection (*26*). Although point mutations in the MHC-I-restricted viral epitopes could subvert CD8^+^ T cell surveillance (*27*), the majority of SARS-CoV-2 T cell epitopes recognized by convalescent individuals or vaccinees immunized with licensed mRNA vaccines, appear unaffected by unique VOC mutations (*28*). These findings indicate a role of cellular immunity in defense against SARS-CoV-2. Here, we observed protection against disease upon challenge infection with VOC B.1.351 after CVnCoV vaccination, despite reduced virus neutralizing titers. These observations suggest that either complementary immune mechanisms are effective or that residual virus neutralizing titers against B.1.351 are sufficient for *in vivo* neutralization in this model. In line with this, it has recently been demonstrated in a non-human primate infection model that relatively low neutralizing antibody titers can protect from SARS-CoV-2-related clinical signs (*26*). The precise contribution of various immune compartments to CVnCoV efficacy requires further evaluation.

Our proof-of-principle study demonstrates that an mRNA vaccine can protect hACE2 mice against disease caused by SARS-CoV-2 independent from the lineages or variant SARS-CoV-2.

## Acknowledgments

We acknowledge Mareen Lange, Silvia Schuparis, Patrick Zitzow and Laura Timm for their excellent technical assistance and Frank Klipp, Doreen Fiedler, Christian Lipinski, Bärbel Berger, and Kerstin Kerstel for their invaluable support in the animal facility. We are very thankful to Thorsten Wolf for providing the SARS-CoV-2 B.1.351 isolate “NW-RKI-I-0028”, and to Roman Wölfel for providing SARS-CoV-2 isolate “BavPat1”.

## Author’s contribution

Conceptualization: DH, BC, SR, JM, SOM, TCM, BP, AD, MB. Methodology: DH, BC, SR, NR, JM, SOM, CW, AB, KW, BP, AD, MB. Formal analysis: DH, BC, NJH, LU, JS, AK, AB, CW. Investigation: DH, BC, NJH, LU, CF, JS, AK, AB, AM, FS, CW. Resources: SR, NR, JM, TCM, SOM, BP, AD, MB. Data Curation: DH, BC, NJH, LU, JS, CW, AD, MB. Writing – original draft preparation: DH, BC, NJH, LU, CW, AD, MB. Writing – review and editing: DH, BC, SR, NR, AB, KW, CW, SOM, TCM, BP, AD, MB. Visualization: DH, BC, NJH, LU, CF, CW. Supervision: DH, BC, AB, KW, SOM, TCM, BP, AD, MB. Project administration: DH, BC, SR, NR, JM, SOM, TCM, BP, AD, MB. Funding acquisition: SR, SOM, BP, AD, MB, TCM.

