## Supplemental Data for "CVnCoV protects human ACE2 transgenic mice from ancestral B BavPat1 and emerging B.1.351 SARS-CoV-2"

### Material and Methods

#### Ethics

The animal experiments were evaluated and approved by the ethics committee of the State Office of Agriculture, Food safety, and Fishery in Mecklenburg – Western Pomerania (LALLF M-V: LVL MV/TSD/7221.3-1-055/20). All procedures using SARS-CoV-2 were carried out in approved biosafety level 3 (BSL3) facilities.

#### Vaccination

Before challenge with SARS-CoV-2 mice were vaccinated prime day 0 and boost day 28 with either NaCl (SHAM), formaldehyde inactivated whole virus preparation (FI-Virus) or an mRNA vaccine (CVnCoV) (Table S1).

CVnCoV is based on the RNActive® platform (claimed and described in e.g. WO2002098443 and WO2012019780) and is comprised of a 5′ cap structure, a GC-enriched open reading frame (ORF), 3′ UTR, polyA tail and does not include chemically modified nucleosides. Lipid nanoparticle (LNP)-encapsulation of mRNA was performed by Acuitas Therapeutics (Vancouver, Canada). The LNPs used in this study are particles of ionizable amino lipid, phospholipid, cholesterol and a PEGylated lipid. The mRNA encoded protein is based on the spike glycoprotein of SARS-CoV-2 NCBI Reference Sequence NC_045512.2, GenBank accession number YP_009724390.1 and encodes for full length S featuring K986P and V987P mutations. For this study we used 8 µg of CVAC20-64_NIAR_R9515 RNActive® (CVnCoV).

For comparison to CVnCoV, we used FI-Virus combined with Alhydrogel® adjuvant. For this, 200mL SARS-CoV-2 Germany/BavPat1/2020 (details see section challenge infection) supernatant was concentrated using PEG Virus Precipitation Kit (Biovision # BIV-K904) to a volume of 2mL. Afterwards this preparation was inactivated by formaldehyde (37%) at a dilution of 1:2000 at 37°C for 24 h. Inactivation of the virus was confirmed by inoculation of

VeroE6 cells. When no cytopathogenic effect (CPE) was detected, cell supernatants were passaged for three passages. For vaccination, freshly prepared stocks of $10^6$ tissue infectious dose 50 ($TCID_{50}$) FI-Virus were mixed with 2% (final concentration) of Alhydrogel® in PBS. Prior vaccination, 20µl of NaCl (SHAM control), FI-Virus or CVnCoV were loaded into single-use insulin syringe with an integrated needle (30G) no longer than 2h before injection. First, the mice were anesthetized by inhalation of isoflurane and the hind leg was shorn with an electric clipper. For all groups 20 µL of the preparation was administered intramuscularly (i.m.) into the M. tibialis (day 0 right leg or left day 28). Before animals were placed back into their cages, 100-140µl blood samples were obtained by puncture of the V. facialis on day 0, day 28, and day 55. For blood collection, the animals remained anesthetized under isoflurane anesthesia (5 vol. %). All groups were monitored for side effects of the injection, and were scored at 24 h post injection. The injection sides were slightly swollen in all groups 24h post vaccination, which resolved after 48-72h.

Serum collection

All blood samples were collected into Z-clot activator 200 µl microtube (Sarstedt). The samples were incubated at room temperature (RT) for 0.5-1h and afterwards centrifuged for 5 min, 10,000 rcf, at RT. All serum samples were stored at <-70°C.

Virus preparation

SARS-CoV-2 Germany/BavPat1/2020 (BavPat1) (GISAID accession EPI_ISL_406862) was kindly provided by Bundeswehr Institute of Microbiology, Munich, Germany. SARS-CoV-2 hCoV-19/Germany/NW-RKI-I-0029/2020 B.1.351-linage or VOC 202012/02 (B.1.351) (GISAID accession EPI_ISL_803957) was kindly provided by Robert-Koch-Institut, Berlin, Germany. Virus stocks were propagated (three passages two passages, respectively) on Vero E6 cells (Collection of Cell Lines in Veterinary Medicine CCLV-RIE 0929) using a mixture of equal volumes of Eagle MEM (Hanks' balanced salts solution) and Eagle MEM (Earle's balanced salts solution) supplemented with 2 mM L-Glutamine, nonessential amino acids adjusted to 850 mg/L, NaHCO3, 120 mg/L sodium pyruvate, 10% fetal bovine serum (FBS), pH 7.2. The virus was harvested after 72h, titrated on Vero E6 cells and stored at -80°C until further use.

Sequencing of the viral genome

Full genome sequencing of SARS-CoV-2 B1.351 hCoV-19/Germany/NW-RKI-I-0029/2020 P2+1 (passage of hCoV-19/Germany/NW-RKI-I-0029/2020) was performed using high-throughput sequencing (HTS). For this purpose, RNA was extracted from cell culture supernatant using a combined TRIzol LS Reagent (Invitrogen, Waltham, MA, USA) and QIAamp RNeasy Mini Kit (Qiagen, Hilden, Germany) protocol. The resulting RNA extracts were subjected to library preparation as detailed described (*1*). The resulting library L4550 was quality-checked, quantified and sequenced on the Ion Torrent S5XL platform on an Ion 530 sequencing chip using 400 bp chemistry.

The Genome Sequencer software suite (versions 2.6; Roche) was applied to execute reference mapping analyses. Since no complete whole-genome sequence was available of the original isolate (hCoV-19/Germany/NW-RKI-I-0029/2020|EPI_ISL_803957|2020-12-28), the genome sequence of SARS-CoV-2 B.1.351 isolate hCoV-19/Germany/BW-ChVir22275/2021|EPI_ISL_875344|2021-01-15 was used as initial reference. Subsequently, the mapping analysis was repeated with the obtained SARS-CoV-2 genome sequence as reference and the corresponding dataset to compile the final SARS-CoV-2 genome sequence of the sample. This determined whole genome sequence of sample L4550 was set as reference for variant calling. The Torrent Suite plugin Torrent variantCaller (version 5.12) was used to detect single nucleotide polymorphism (SNP) variants (parameter settings: generic, S5/S5XL(530/540), somatic, low stringency, changed alignment arguments for the TMAP module from map 4 [default] to map1 map2). Identified SNP variants were visualized with

Geneious Prime (10.2.3; Biomatters, Auckland, New Zealand) and compared with the SNP variants detected using the variant analysis tool implemented in Geneious Prime (default settings, minimum variant frequency 0.02).

The SARS-CoV-2 genome sequence generated in this study is available under the ENA Study accession number PRJEB43810.

Challenge infection

Mice in groups of up to five animals were kept in individually ventilated cages (IVCs) for the entire study (Table S1). The animals were infected under short-term isoflurane inhalation anesthesia with 25 µl of either $10^{5,875}$ TCID50 SARS-CoV-2 BavPat1 (calculated from back-titration of the original material) or $10^{5,5}$ TCID50 SARS-CoV-2 B1.351 (calculated from backtitration of the original material) per animal. To prevent spill-over between different pairs, the IVCs were strictly separated in individual cage systems. During the entire study, all animals were offered water ad libitum, and were fed and checked for clinical scores and body weight daily by animal caretakers and study researchers. A nasal swab sample of each animal was taken at 4dpi under short-term isoflurane inhalation anesthesia. Animals with signs of severe clinical symptoms and/or body weight loss over 20% were euthanized before the endof the study. All animals were euthanized at day 10 post infection.

RNA extraction and RT-qPCR

RNA from combined nasal/buccal swabs and organ samples was extracted using the NucleoMag® VETkit (Macherey-Nagel, Düren, Germany) in combination with a Biosprint 96 platform (Qiagen, Hilden, Germany). Each extracted sample was eluted in 100µl. Viral RNA genome was detected and quantified by real-time reverse transcription polymerase chain reaction (real-time RT-qPCR) on a BioRad real-time CFX96 detection system (BioRad, Hercules, USA). Target sequence for amplification was the viral RNA-dependent RNA polymerase (WHO. https://www.who.int/docs/default-source/coronaviruse/real-time-rt-pcr-assays-for-the-detection-of-sars-cov-2-institut-pasteur-paris.pdf?sfvrsn=3662fcb6_2). Genome copies per µl RNA template were calculated based on a quantified standard RNA, where absolute quantification was done by the QX200 Droplet Digital PCR System in combination with the 1-Step RT-ddPCR Advanced Kit for Probes (BioRad, Hercules, USA).

RBD antibody Elisa

Sera were analysed using ELISA performed as previously described (*2*). Briefly, Elisa plates (Greiner Bio-One GmbH) were coated with 100 ng/well the RBD overnight at 4°C in 0.1 M carbonate buffer (1.59 g Na2CO3 and 2.93 g NaHCO3, ad. 1 L aqua dest., pH 9.6) or were treated with the coating buffer only. Afterwards, the plates were blocked for 1 hr at 37°C using 5% skim milk in phosphate-buffered saline (PBS). Sera were pre-diluted 1/100 in TBS-Tween (TBST) and incubated on the coated and uncoated wells for 1 hr at RT. A multi-species conjugate (SBVMILK; obtained from ID Screen® Schmallenberg virus Milk Indirect ELISA; IDvet) was diluted 1 1/80 and then added for 1 hr at RT. Following the addition of tetramethylbenzidine (TMB) substrate (IDEXX), the ELISA readings were taken at a wavelength of 450 nm on a Tecan Spectra Mini instrument (Tecan Group Ltd). Between each step, the plates were washed three times with TBST. The adsorbance was calculated by subtracting the optical density (OD) measured on the uncoated wells from the values obtained from the protein-coated wells for the respective sample.

Virus neutralization test (VNT)

To evaluate specifically the presence of virus neutralizing antibodies in serum samples, we performed a VNT. Therefore, sera were pre-diluted 1/16 or 1/32 -with DMEM in a 96 well deep well master plate. Three times 100 µl, representing three technical replicates, of this pre-dilution were transferred into a 96 well plate. A log2 dilution was conducted by passaging 50 µl of the serum dilution in 50 µl DMEM, leaving 50 µl of sera dilution in each well. Subsequently 50 µl of the respective SARS-CoV-2 (BavPat1 or B.1.351) virus dilution (100 TCID50/well) was added to each well and incubated for 1 hour at 37 °C. Lastly, 100 µl of trypsinated VeroE6 cells (cells of one confluent TC175 flask per 100 ml) in DMEM with 1 % penicillin/streptomycin supplementation was added to each well. After 72 hours incubation at 37 °C, the wells were evaluated by light microscopy. A serum dilution was counted as neutralizing in the case no specific CPE was visible. The virus titer was confirmed by virus titration, positive and negative serum samples were included.

### Supplementary Figure legends

**Fig.S1. Experimental design.** K18-hACE2 mice were either vaccinated at day 0 (prime) and day 28 (boost) i.m. with 20µl of 8µg CVnCoV, or received 20µl NaCl (Sham) or 20µl of $10^6$ $TCID_{50}$ FI-Virus+2%Alhydrogel® in PBS which served as control groups. Blood samples were collected at day 0, day 28 and day 55. Mice were either challenged with $10^5$ SARS-CoV-2 BavPat1 (Sham n=4, FI-Virus n=10, CVnCoV n=10) or $10^5$ B.1.351 (Sham n=5, FI-Virus n=10, CVnCoV n=10) at day 59 for a total of 10 days. An oral swab was taken at day 4. Viral load was determined in selected organs at day 10 or when animals reached the humane endpoint of the study. Image was generated with Biorender (Biorender.com)

**Fig.S2 Cell culture passage of the VOC B.1.351 NW-RKI-I-0028 strain did not alter genome sequence of the challenge virus stock L4550.** All characteristic mutations of VOC B.1.351 (upper panel) were detected in the deeply sequenced stock of the challenge virus B.1.351 NW-RKI-I-0028 (L4550) as marked by asterisks. Additional alterations for the B.1.351 NW-RKI-I-0028 and NW-RKI-I-0029 strains and L4550 cell culture passage are shown (lower panel). Mean coverage of the virus genome sequence of L4550 was 13,184 (± 7,958.4) reads. No single nucleotide polymorphism variants were detected in the mutations (L18F, D80A, D215G, deletion De241-243, K417N, E484K, N501Y, A701V) and in the cleavage site of the spike glycoprotein.

**Fig. S3 CVnCoV protects K18-hACE2 mice against SARS-CoV-2 variants BavPat1 and B.1.351.** K18-hACE2 mice were vaccinated and challenged with SARS-CoV-2 variants BavPat1 or B.1.351 as indicated in Fig. 1. (**A, C** and **E**) Body weight and (**B, D** and **F**) clinical score for K18-hACE2 mice in the (**A** and **B**) SHAM, (**C** and **D**) FI-Virus and (**E** and **F**) CVnCoV groups was monitored daily. Lines represent individual animals over the course of the experiment (BavPat1 = green; B.1.351 = red).

Table S1: Experimental and control groups.

| group | No. of animals | Vaccine | Dose, volume, route | Dosing | Serum collection | Challenge infection | | End of the study |
|---|---|---|---|---|---|---|---|---|
| I | K18-hACE2 mice Female n=10 | CVnCoV | 8 µg, 20 µl, i.m. | d0, 28 | d0, d28, d55 | BavPat1 | | d69 (10dpc) |
| II | | | 8 µg, 20 µl, i.m. | | | | B1.351 | |
| III | | Formalin-inactivated virus +Alum | $10^{6}$ $TCID_{50}$, 20µl, i.m. | | | BavPat1 | | |
| IV | | | $10^{6}$ $TCID_{50}$, 20µl, i.m | | | | B1.351 | |
| V | n=5 | Sham (NaCl) | -, 20 µl, i.m. | | | BavPat1 | | |
| VI | | | -, 20 µl, i.m. | | | | B1.351 | |

Table S2. Sequencing coverage of L4550 for mutations typically found in the spike protein gene of B.1.351 variants and the spike protein polybasic cleavage site.

| B.1.351 mutations Spike | Coverage | Variant |
|---|---|---|
| **L18F** | 8,584 reads | None |
| **D80A** | 6,361 reads | None |
| **D215G** | 9,898 reads | None |
| **Deletion 241-243** | (9628 reads) | None |
| **K417N** | 8,606 reads | None |
| **E484K** | 9,094 reads | None |
| **N501Y** | 9,868 reads | None |
| **A701V** | 14,038 reads | None |
| **Cleavage site** | 17,730 reads | None |

Fig. S1

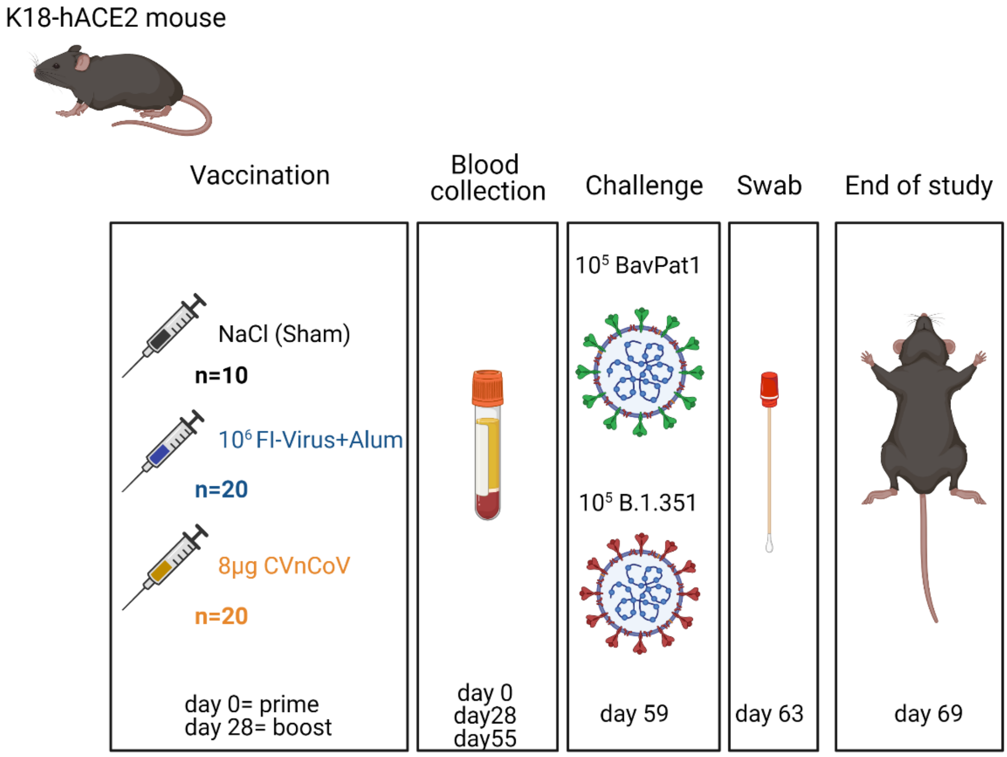
K18-hACE2 mouse
Vaccination
Blood collection
Challenge
Swab
End of study
NaCl (Sham)
n=10
$10^6$ FI-Virus+Alum
n=20
8µg CVnCoV
n=20
day 0= prime
day 28= boost
day 0
day28
day55
$10^5$ BavPat1
$10^5$ B.1.351
day 59
day 63
day 69

Fig. S2

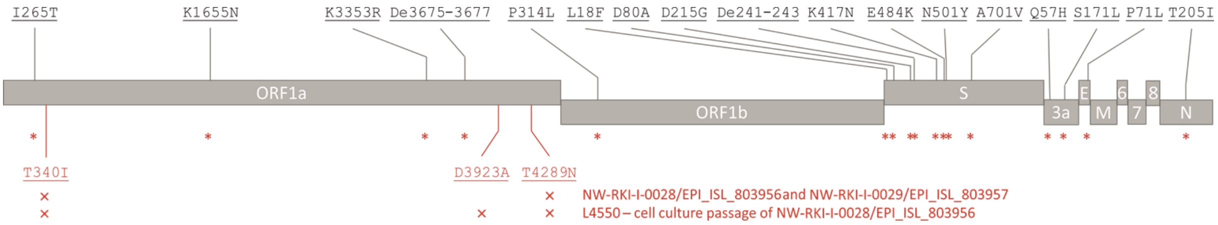
I265T
K1655N
K3353R
De3675-3677
P314L
L18F
D80A
D215G
De241-243
K417N
E484K
N501Y
A701V
Q57H
S171L
P71L
T205I
ORF1a
ORF1b
S
3a
E
M
6
7
8
N
T340I
D3923A
T4289N
NW-RKI-I-0028/EPI_ISL_803956and NW-RKI-I-0029/EPI_ISL_803957
L4550 – cell culture passage of NW-RKI-I-0028/EPI_ISL_803956

Fig. S3

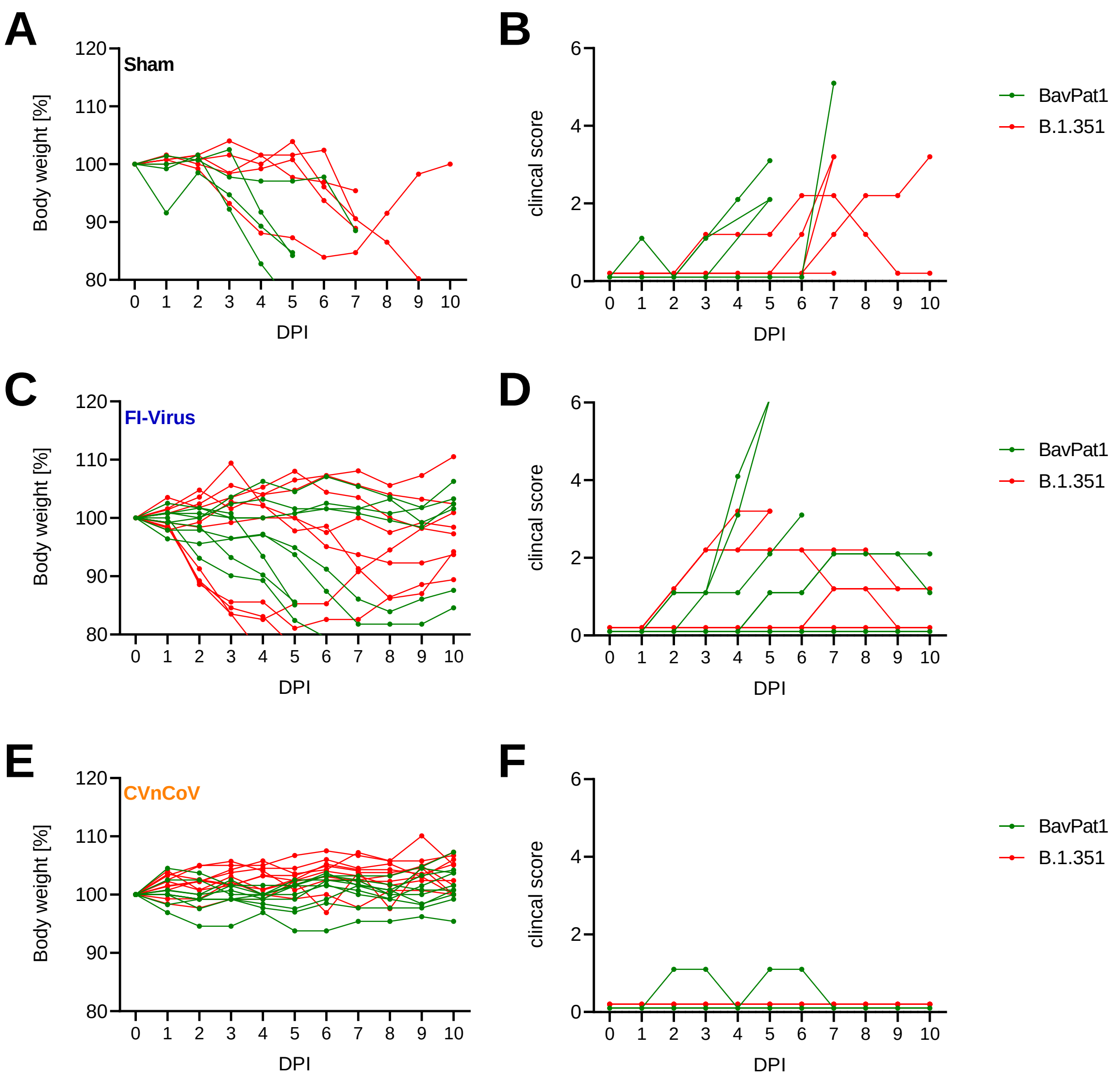

A
Sham
Body weight [%]
DPI
B
clincal score
DPI
BavPat1
B.1.351
C
FI-Virus
Body weight [%]
DPI
D
clincal score
DPI
BavPat1
B.1.351
E
CVnCoV
Body weight [%]
DPI
F
clincal score
DPI
BavPat1
B.1.351
